# MutBERT: Probabilistic Genome Representation Improves Genomics Foundation Models

**DOI:** 10.1101/2025.01.23.634452

**Authors:** Weicai Long, Houcheng Su, Jiaqi Xiong, Yanlin Zhang

## Abstract

**Motivation:** Understanding the genomic foundation of human diversity and disease requires models that effectively capture sequence variation, such as single nucleotide polymorphisms (SNPs). While recent genomic foundation models have scaled to larger datasets and multi-species inputs, they often fail to account for the sparsity and redundancy inherent in human population data, such as those in the 1000 Genomes Project. SNPs are rare in humans, and current masked language models (MLMs) trained directly on whole-genome sequences may struggle to efficiently learn these variations. Additionally, training on the entire dataset without prioritizing regions of genetic variation results in inefficiencies and negligible gains in performance.

**Results:** We present MutBERT, a probabilistic genome-based masked language model that efficiently utilizes SNP information from population-scale genomic data. By representing the entire genome as a probabilistic distribution over observed allele frequencies, MutBERT focuses on informative genomic variations while maintaining computational efficiency. We evaluated MutBERT against DNABERT-2, various versions of Nucleotide Transformer, and modified versions of MutBERT across multiple downstream prediction tasks. MutBERT consistently ranked as one of the top-performing models, demonstrating that this novel representation strategy enables better utilization of biobank-scale genomic data in building pretrained genomic foundation models.

**Availability:** https://github.com/ai4nucleome/mutBERT

## Introduction

Recent advancements in natural language processing (NLP) have driven transformative progress in genomics, particularly through the development of genomic foundation models (Benegas et al., 2025; Consens et al., 2023). These models leverage deep learning techniques to analyze DNA sequences, enabling the prediction of functional elements, regulatory mechanisms, and variant effects. Building on the success of models like BERT (Devlin, 2018) and GPT (Radford, 2018) in NLP, the genomics community has adapted these concepts to address challenges such as modeling long-range dependencies in DNA sequences (Nguyen et al., 2024), (Cheng et al., 2025) and improving predictions based on genomic data (Benegas et al., 2023).

The development of DNA foundation models began in 2021 with DNABERT (Ji et al., 2021), which utilized overlapping k-mers (Celikkanat et al., 2024) as tokens and the original BERT architecture to analyze DNA sequences. Trained exclusively on the human reference genome, DNABERT could handle sequences up to 500 base pairs and established an early standard for genomic representation learning. However, its tokenization methods, constrained sequence lengths, and an outdated architecture restricted its ability to generalize to broader and more complex genomic contexts.

Building on these foundations, the rapid progress in NLP during subsequent years, including the rise of ChatGPT (Achiam et al., 2023), introduced advanced methodologies and architectures that were adapted to genomic modeling (Sanabria et al., 2024). For instance, DNABERT-2 (Zhou et al., 2023) incorporated byte pair encoding (BPE) tokenization (Sennrich, 2015) to improve data representation efficiency. It also adopted ALiBi (Press et al., 2021) for managing longer sequences and flash attention (Dao et al., 2022) for computational optimization, enabling the model to process multi-species datasets. These innovations, although derived from NLP, demonstrated the potential to extend the capabilities of genomic modeling by enhancing sequence length scalability and data diversity.

Nucleotide Transformer (Dalla-Torre et al., 2024) further explored the benefits of scaling model size and input diversity by training various models on the human reference genome, individual genomes from the 1000 Genomes Project (Byrska-Bishop et al., 2022), and multi-species data. It highlighted the advantages of leveraging both larger models and richer datasets. Meanwhile, models such as HyenaDNA (Nguyen et al., 2024) and MambaDNA (Schiff et al., 2024) addressed the limitations of handling long-range genomic dependencies by extending input lengths to hundreds of kilobases using attention-free architectures. These models successfully overcame the quadratic scaling bottlenecks of traditional transformers, demonstrating their efficacy for tasks requiring large genomic contexts.

Other approaches explored integrating additional data, such as evolutionary information through multispecies sequence alignments. For instance, GPN-MSA (Benegas et al., 2024) introduced a framework that combined target sequences with their corresponding multiple sequence alignments from diverse species to improve variant effect predictions, particularly in functional genomic regions.

These developments reflect three key trends in genomic foundation modeling: increasing model size often enhances performance, innovations like flash attention and ALiBi enable longer sequence contexts, and incorporating diverse datasets—particularly multi-species data—yields more robust models. Despite these advancements, efficiently leveraging data from the same species, such as the 1000 Genomes Project, remains a significant challenge. The 1000 Genomes dataset, though invaluable for understanding human genetic diversity, includes substantial redundancy in DNA segments across individuals, which poses challenges for its effective utilization by conventional methods. Extending these approaches to larger, more complex biobank-scale datasets poses an even greater challenge.

To address these issues, we present MutBERT, a probabilistic genome-based masked language model designed to efficiently utilize SNP information from population-scale genomic data. Unlike previous approaches, MutBERT represents genomic sequences as probabilistic distributions over allele frequencies, enabling it to focus on informative regions while mitigating redundancy. Our results show that MutBERT performs well on the 1000 Genomes dataset, effectively capturing population-level variation. Furthermore, we believe that MutBERT’s probabilistic representation strategy positions it to better utilize biobank-scale datasets in the future.

By consistently ranking among the top-performing models across multiple downstream tasks, MutBERT demonstrates its potential as a scalable, biologically informed genomic foundation model. This work provides insight into novel strategies for in leveraging population-scale genomic datasets to advance the field of genomics.

## Materials and methods

### Model Architecture

We proposed MutBERT and its two variants, MutBERT-Ref and MutBERT-Multi, all sharing the same network architecture. MutBERT and MutBERT-Multi utilize probabilistic genome representations as input for masked language modeling (MLM). These probabilistic genomes are derived from the 1000 Genomes Project and 100-way multispecies alignments (Benegas et al., 2024), respectively. In contrast, MutBERT-Ref takes one-hot embeddings of the reference genome as input, providing a deterministic genome baseline for comparison.

As shown in Fig. 1, MutBERT is based on the Transformer Encoder architecture (Vaswani, 2017), incorporating recent advancements (Su et al., 2024), (Dao et al., 2022), (Liu et al., 2023) in deep learning to enhance efficiency and scalability for genomic data. MutBERT comprises 12 encoder layers, with GELU (Dauphin et al., 2017) as the activation function for the feed- forward network. It has 86 million learnable parameters. Unlike other models that take a sequence of tokens as input, MutBERT processes a probabilistic matrix representation. Due to this unique input format, traditional embedding layers implemented in widely used deep learning packages such as PyTorch (Paszke et al., 2019) are not directly applicable. Instead, we employ a linear layer to map the probabilistic input matrix into *d*-dimensional embeddings, ensuring compatibility with the Transformer Encoder architecture. Briefly, traditional embedding layers in language models typically function as look-up tables, assigning each discrete token a corresponding embedding vector. This process can be implemented as a matrix multiplication between a one-hot encoded vector and an embedding matrix, effectively selecting the embedding associated with a single token. In contrast, our probabilistic genomic representation assigns each locus a distribution over multiple possible states (e.g., A, T, C, G). Thus, by assigning each state an embedding vector, the final embedding at a given locus is computed as a weighted sum over all embeddings, with weights corresponding to the probabilities of each state. Consequently, we use a linear layer to transform our probabilistic genomic representation into the appropriate embedding space. To address the limitations of fixed input lengths in traditional positional embeddings, we integrate Rotary Position Embedding (RoPE) (Su et al., 2024), which enables effective handling of variable-length inputs and ensures robust performance across diverse genomic contexts. Additionally, we utilize Flash Attention (Dao et al., 2022) to optimize computation and memory efficiency, significantly reducing the quadratic scaling bottleneck associated with standard attention mechanisms, allowing MutBERT to process longer sequences efficiently. MutBERT also includes support for Low-Rank Adaptation (LoRA) (Hu et al., 2021) during the fine-tuning stage, enabling parameter-efficient training by adapting only a small subset of parameters. However, as the model presented in this study is relatively small (86 MB), we did not applied LoRA in our experiments.

**Fig. 1:**
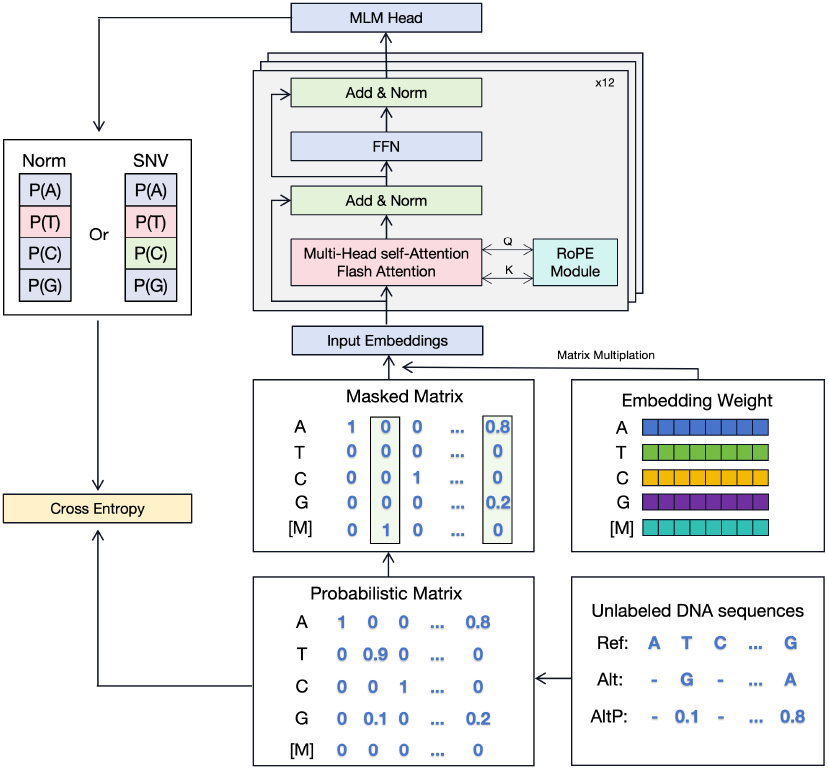
Overview of MutBERT.

### Probabilistic Genome Representation

In the probabilistic genome representation, each nucleotide position is encoded as a probabilistic distribution over the four nucleotide bases: A, T, C, and G, derived from allele frequencies in population-scale datasets such as the 1000 Genomes Project. This approach captures genetic diversity across individuals, providing a nuanced representation of both conserved and variable genomic regions.

This method draws inspiration from established genomic practices, such as using position-specific weight matrices (PSWMs) to represent sequence motifs and inferring ancestral states from pairwise alignments. Similar to how PSWMs capture the likelihood of observing specific bases at each position in a motif, the probabilistic genome representation encodes the likelihood of each nucleotide at every position based on observed allele frequencies. Positions without observed variants are represented deterministically, with the reference nucleotide assigned a probability of 1, while variable positions are represented as distributions reflecting the observed allele frequencies.

Under this formulation, a DNA sequence of length *L* can be represented as a probabilistic matrix of size 4 *× L*, where each column corresponds to the probability distribution of the four nucleotides at a specific genomic position. Unlike existing models that rely solely on a reference genome or individual DNA sequences as input, our probabilistic genomic representation integrates additional genomic variation data to construct these nucleotide probability distributions. Specifically, these distributions can be derived either by computing allele frequencies from population-scale genomic datasets or by extracting variant frequencies from multiple-sequence alignment data available through resources such as the UCSC Genome Browser (Blanchette et al., 2004). In subsequent sections, we detail how we generated these probabilistic inputs to train MutBERT, and MutBERT-Multi models.

In the development of MutBERT, we extend it to account for five special tokens, such as [PAD], [CLS], and [MASK], which expand the matrix representation to 9 *× L*. This expanded format ensures compatibility with masked language modeling while preserving the probabilistic representation of the underlying genomic data.

### Tokenization and Vocabulary

To ensure high-quality input data, we discard DNA sequences containing ambiguous nucleotides represented by N. The input to our model is processed at the individual nucleotide level, resulting in a vocabulary consisting of A, T, C, and G. Additionally, our vocabulary includes the special tokens [PAD], [CLS], [SEP], [UNK], and [MASK], commonly used in masked language modeling.

### Pretraining via Masked Language Modeling

#### Mask Strategy

We adopt a masked language modeling (MLM) setup similar to that used in BERT, employing a masking ratio of 15%. To prioritize biologically relevant information, we focus on masking single nucleotide variant (SNV) tokens. If the number of SNV tokens exceeds 15% of the sample length, we randomly select 15% of the tokens. Otherwise, we supplement the selection with non-SNV tokens to achieve the required masking ratio. For the selected tokens, we replace them with a [MASK] token 80% of the time and leave them unchanged 20% of the time. Since MutBERT operates on a probabilistic matrix as input, we apply masking by modifying the probability distribution of the selected tokens. Specifically, we assign a probability of 1 to the [MASK] token in the masked position, ensuring compatibility with the probabilistic input format.

#### Loss Function

The dataset includes mutation probabilities as part of the input representation, which we use as target labels during pretraining. We mask the SNV tokens and train the model to predict these probabilities. The loss is computed as the cross-entropy (Mao et al., 2023) between the predicted logits and the nucleotide probabilities as follows:

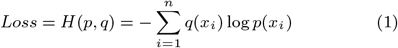

Where *p*_(_*x*_*i*_) represents the predicted probability for label *x*_*i*_, and *q*_(_*x*_*i*_) represents true probability of *x*_*i*_. Since *q*(*x*_*i*_) corresponds to nucleotide probabilities at locus *i*, these probabilities are often extremely small or very close to 1, causing the training loss to insufficiently guide the model in learning effectively from such highly skewed distributions. Inspired by the concept of temperature scaling from knowledge distillation (Hinton et al., 2015), we propose to modify *q*(*x*_*i*_) using temperature scaling as follows:

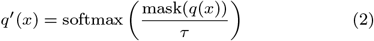

where mask selectively filters out the non-zero elements of *q*(*x*), and *τ* denotes the temperature parameter. We then employed softmax to normalize *q*^*′*^(*x*_*i*_). To evaluate the impact of temperature scaling, we trained models with no scaling (*τ* = 1) as well as with temperatures of *τ* = 0.7 and *τ* = 2. Among these, the model trained with *τ* = 0.7 consistently showed superior performance. Therefore, all results reported in this paper are based on *τ* = 0.7.

### Models and Pre-training Data

We trained three models, each with the same architecture and training procedure. The key difference among these models lies in the source of their training data. The datasets used for training are summarized in Table 1 and include the 1000 Genomes Project data, multispecies data, and the human reference genome. For all models, we performed training on all autosomes and the X chromosome. In each dataset, we removed all N characters locus in the human reference genome and concatenated the remaining DNA segments using the [SEP] token. Among all settings, We used chromosome 22 as the validation set and the remaining chromosomes as the training set.

**Table 1.** Pretraining dataset statistics for Human-Ref, 1000G, and Multispecies. Each sample represents a 510 bp DNA sequence. The number of samples refers to the total number of DNA sequences randomly sampled from the genome.

| Pretraining dataset<br>split | Human-Ref |  | 1000G |  | Multi |  |
| --- | --- | --- | --- | --- | --- | --- |
|  | Train (M) | Test (M) | Train (M) | Test (M) | Train (M) | Test (M) |
| Total bp | 2872 | 39 | 2872 | 39 | 2872 | 39 |
| Num. SNVs | 0 | 0 | 9(0.34%) | 0.1(0.37%) | 2008(69.92%) | 25(64.60%) |
| Num. Samples | 30 | 0.15 | 30 | 0.15 | 30 | 0.15 |
| Num. Samples w/ SNVs | 0 | 0 | 21(71.49%) | 0.11(72.06%) | 29(96.90%) | 0.13(92.68%) |

All models were trained using a batch size of 2048 and a maximum sequence length of 512. The training process consisted of 120,000 steps, employing the AdamW optimizer with parameters *β*_1_ = 0.9, *β*_2_ = 0.99, *ϵ* = 1*e −* 8 and weight decay of 0.01. During the initial 10,000 steps, the learning rate was linearly increased from 0 to 4*e −* 4. For the remaining 110,000 steps, the learning rate decayed gradually following a cosine schedule, reaching 1e-8 at the final step.

#### MutBERT

It was trained on probabilistic genome data derived from the 1000 Genomes Project. We downloaded Variant Calling Format (VCF) files from the 1000 Genomes Project, which includes data from 3,202 high-coverage genomes spanning 27 geographically structured populations. For each genomic locus, we calculated allele frequencies from the SNP information provided in the VCF files, discarding rare mutations with allele frequencies less than 2%. Using these allele frequencies, we created probabilistic training data, assigning probabilities to each nucleotide at every position based on their observed frequencies. This approach provided a probabilistic representation of human genetic diversity for training MutBERT.

#### MutBERT-Multi

It was trained on probabilistic genomes derived from the multiz whole-genome alignment dataset (Blanchette et al., 2004). This alignment is anchored to the human genome, with mutation probabilities corresponding to positions in the human genome. Following the approach outlined in GPN-MSA (Benegas et al., 2024), we included 10 closely related primates to humans, resulting in a dataset containing 100 vertebrate species. Mutation probabilities were calculated by counting the number of nucleotides at each position across the alignment. This dataset enabled us to represent multispecies genomic diversity probabilistically, providing a distinct training context for MutBERT-Multi.

#### MutBERT-Ref

It was trained exclusively on data derived from the human reference genome (hg38). The training dataset was generated by retaining all autosomal and X chromosomes. All N characters were removed, and the remaining DNA segments were concatenated using the [SEP] token. Tokenization was performed at the individual nucleotide level, as described previously. This model is similar to other genomic models, such as DNABERT and Nucleotide Transformer trained on the human reference genome, but it employs single nucleotide-level tokenization instead of k-mer-based or BPE approaches. To enable MutBERT to train effectively on this deterministic data, we represented the input as a probabilistic matrix, assigning a probability of 1 to the reference allele at all positions.

### Alternative Models

We compared MutBERT against DNABERT-2 (Zhou et al., 2023) and Nucleotide Transformer (NT) (Dalla-Torre et al., 2024). DNABERT-2 is pre-trained on the human genome and 134 other species, incorporating advanced techniques such as BPE, ALiBi, and Flash Attention. These techniques enable DNABERT- 2 to handle sequences longer than those encountered during pretraining, making it highly versatile for various genomic tasks. Nucleotide Transformer represents a family of transformer-based models for DNA sequence analysis. We evaluated three variants of NT-v1: NT-500M-humanRef, NT-500M-1000G, and NT-2500M- multi, pre-trained on the human reference genome, the 1000 Genomes Project, and genomes from 850 species, respectively. NT- v1 models can process sequences up to 1,000 tokens, equivalent to 6,000 base pairs. Additionally, we evaluated NT-v2, which is pre-trained exclusively on the multi-species dataset. NT-v2 includes four variants: NTv2-50M-multi, NTv2-100M-multi, NTv2-250M- multi, and NTv2-500M-multi, with the capability to process sequences up to 2,048 tokens, equivalent to 12,000 base pairs. We utilized pre-trained weights for these models, obtained from Hugging Face, for all evaluations. It is important to note that some NT models are significantly larger than MutBERT. Due to limited computational resources, we applied standard fine-tuning techniques only to models with fewer than 300 million parameters. For larger models, we utilized LoRA for parameter-efficient fine-tuning.

### Downstream Tasks

We developed MutBERT to better utilize large-scale genomic data from individuals of the same species, with a primary focus on training a model for humans. Consequently, we evaluated MutBERT’s performance on tasks specifically designed for human genomic analysis.

#### Transcription factor binding site prediction

We evaluated MutBERT and other foundation models on the task of transcription factor binding site (TFBS) prediction in humans, using datasets from the GUE benchmark (Zhou et al., 2023). This task includes five datasets, each designed as a binary classification task, with input sequences fixed at a length of 100 bp.

#### Nucleotide Transformer downstream tasks

We also assessed MutBERT and other foundation models on 18 datasets from the Nucleotide Transformer benchmark (Dalla-Torre et al., 2024). We split each dataset into training and testing sequences following the established practice in Dalla-Torre et al. The number of sequences used for training and testing, the number of target classes, and the sequence length for each task are shown in Table 2. The benchmark encompasses four primary tasks: Epigenetic Marks Prediction (EMP), Promoter Sequence Prediction (PSP), Enhancer Sequence Prediction (ESP), and Splice Site Prediction (SSP). Chromosomes 21 and 22 were reserved for testing, while the remaining chromosomes were used for fine-tuning. Most datasets involve binary classification tasks, with the exception of splice sites all and enhancers types, which are multi-class classification tasks.

**Table 2.** Detail of NT-downstream tasks

| Task | Num. train seqs. | Num. test seqs. | labels | length |
| --- | --- | --- | --- | --- |
| promoter_all | 30,000 | 1,584 | 2 | 300 |
| promoter_tata | 5,062 | 212 | 2 | 300 |
| promoter_no_tata | 30,000 | 1,372 | 2 | 300 |
| enhancers | 30,000 | 3,000 | 2 | 400 |
| enhancers_types | 30,000 | 3,000 | 3 | 400 |
| splice_sites_all | 30,000 | 3,000 | 3 | 600 |
| splice_sites_acceptor | 30,000 | 3,000 | 2 | 600 |
| splice_sites_donor | 30,000 | 3,000 | 2 | 600 |
| H2AFZ | 30,000 | 3,000 | 2 | 1,000 |
| H3K27ac | 30,000 | 1,616 | 2 | 1,000 |
| H3K27me3 | 30,000 | 3,000 | 2 | 1,000 |
| H3K36me3 | 30,000 | 3,000 | 2 | 1,000 |
| H3K4me1 | 30,000 | 3,000 | 2 | 1,000 |
| H3K4me2 | 30,000 | 2,138 | 2 | 1,000 |
| H3K4me3 | 30,000 | 776 | 2 | 1,000 |
| H3K9ac | 23,274 | 1,004 | 2 | 1,000 |
| H3K9me3 | 27,438 | 850 | 2 | 1,000 |
| H4K20me1 | 30,000 | 2,270 | 2 | 1,000 |

#### eQTL Variant Effect Prediction

The expression quantitative trait loci (eQTL) dataset used in this task is derived from Enformer (Avsec et al., 2021) and Nucleotide Transformer (Dalla-Torre et al., 2024). Each sample in the dataset consists of a reference sequence and an alternative sequence, with the mutation site located at the center. The output is a binary value referring to whether the variant has a causal effect on gene expression.

### Context Length and Speed Optimization

#### Rotary Position Embedding (RoPE)

We pretrained our model at the individual nucleotide level, using input sequences consisting of 512 tokens. However, we intend to fine-tune the pretrained model on sequences of varying lengths, including those significantly shorter or longer than the pretraining length. Therefore, we adopted Rotary Position Embedding (RoPE) (Su et al., 2024), an effective positional encoding approach for transformer-based models. Unlike traditional positional embeddings, RoPE encodes positions through rotations in a complex-valued space, naturally capturing relative positional relationships among tokens. This method improves computational efficiency and enhances the model’s ability to generalize across different sequence lengths.

#### Dynamic Neural Tangent Kernel (NTK) Extrapolation

To extend RoPE for handling longer sequences, a dynamic scaling method was introduced (Liu et al., 2023), where the scaling parameter *α* is adjusted dynamically based on the sequence length. For sequences shorter than or equal to the pretraining context length, the original positional encoding is used. For longer sequences, the scale parameter *α* is modified according to the ratio of the current sequence length to the original pretraining context length, resulting in a progressively increasing scale as the sequence length grows. The scaling of *α* is computed as:

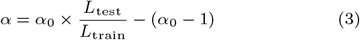

Where *α*_0_ is the initial scaling parameter, *L*_test_ is the sequence length during testing, and *L*_train_ is the sequence length used during training. The updated positional encoding *θ* is then computed as:

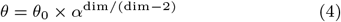

Where *θ*_0_ is the initial rotary parameter, and dim represents the dimensionality of the positional encoding.

#### Flash Attention

Flash Attention is an efficient algorithm designed to perform exact standard attention computations with reduced memory and time overhead. By minimizing IO operations between the GPU’s fast on-chip memory and high-bandwidth memory, Flash Attention enables faster and more memory-efficient attention calculations. This is achieved by splitting Key, Query, and Value matrices into smaller blocks and incrementally computing the softmax operation. We utilized the publicly available Flash Attention implementation in PyTorch (Paszke et al., 2019) to enhance the computational efficiency of MutBERT, allowing it to handle longer input sequences without compromising performance.

#### Low-Rank Adaptation (LoRA)

Let *W*_*o*_, *W ∈* ℝ ^*m×n*^ represent the original weight matrix before fine-tuning and the task-specific fine-tuned weight matrix, respectively. The relationship between them can be expressed as:

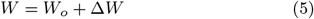

where Δ*W ∈*ℝ ^*m×n*^ captures the task-specific updates to the weights. In conventional fine-tuning, every element in Δ*W* is updated independently, resulting in high computational and memory demands. LoRA addresses this by modeling Δ*W* using a low-rank decomposition, and only updates *A* and *B*:

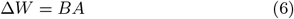

where *B ∈*ℝ ^*m×r*^, *A ∈*ℝ ^*r×n*^, and *r ≪* min(*m, n*). This decomposition reduces the number of trainable parameters from *m × n* to *r ×* (*m* + *n*), substantially lowering the computational and memory requirements.

### Finetune and Sequence Embedding

We used AdamW (Loshchilov, 2017) as the optimizer in all deep learning based finetuning tasks. Fine-tuning was performed for 1,000 steps, including 50 warm-up steps, with a batch size of 32. For MutBERT, DNABERT-2, NTv2-50M-Multi, NTv2- 100M-Multi and NTv2-250M-Multi, we performed standard fine-tuning with a learning rate of 3e-5, while for larger Nucleotide Transformer variants, we applied LoRA with a learning rate of 1e-4, LoRA alpha of 16, LoRA dropout of 0.05, and LoRA r of 8. All models were trained using the Transformers library (Wolf et al., 2020). Following established practice (Zhou et al., 2023), we used Matthews correlation coefficient (MCC) (Jurman et al., 2012) to evaluate model performance on TFBS and Nucleotide Transformer downstream datasets. During the training process, the model was validated every 200 steps. We implemented an early stopping strategy based on the Matthews correlation coefficient (MCC) metric, with a patience of 5 validation checks. Each model was trained using 3 different random seeds, and the average results across these runs are reported.

For the eQTL variant effect prediction task, we used area under the receiver operating characteristic curve (AUROC) as the evaluation metric. Following previous research (Schiff et al., 2024; Kao et al., 2024), we extracted embeddings by averaging a 1,536 base pair window around the SNV positions in the reference and alternative sequences, then concatenating them along the channel dimension. Since the maximum supported sequence length varies by model, we used 12,000 base pairs for Nucleotide Transformer and 2,048 base pairs for DNABERT-2 to obtain embeddings. We also used 2048 base pairs for MutBERT. To achieve this, we extended MutBERT’s sequence length from 512 to 2048. The embeddings were used to train an SVM with an RBF kernel, stratified by distance to the nearest transcription start site (TSS). For each TSS distance bucket, 5,000 training samples were randomly selected, and the AUROC was recorded on the test set. This process was repeated five times, and the average score was reported.

## Results

After pretraining, MutBERT achieved an evaluation loss of 0.691 and a perplexity of 1.996. MutBERT-Ref had an evaluation loss of 0.803 and a perplexity of 2.233, while MutBERT-Multi had an evaluation loss of 1.077 and a perplexity of 2.936. All on test data. To evaluate the effectiveness of our probabilistic genome representation and pretraining method, we compared MutBERT against state-of-the-art genomic language models, including DNABERT-2 and Nucleotide Transformer (NT), both of which represent leading MLM-trained foundation models for genomic data. Additionally, we included MutBERT-Ref and MutBERT-Multi to further dissect the impact of our newly proposed probabilistic genome representation. A summary of the models involved in this study is provided in Table 3. Most of these models contain more parameters and are trained on more tokens than MutBERT.

**Table 3.**
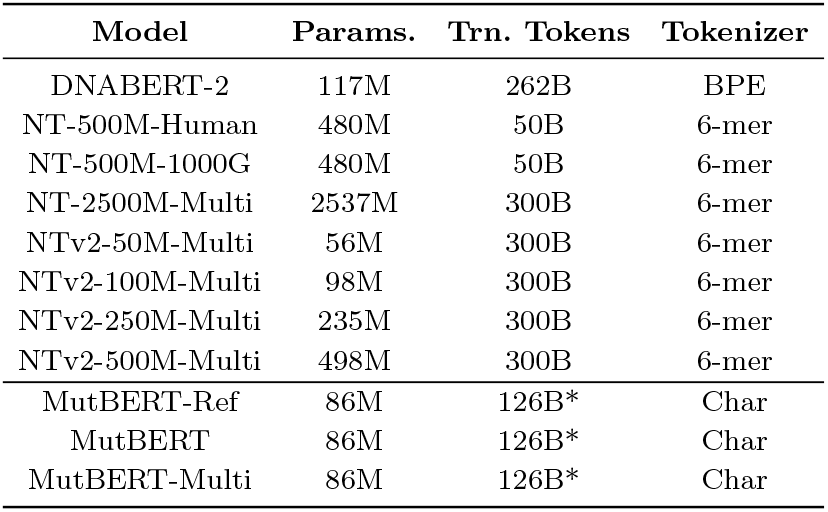
The statistics of each model. The 3 columns represent the number of model parameters, the number of training tokens and the tokenizer used in pre-training. Numbers marked by * indicate the total number of tokens seen throughout 120K training iterations, corresponding to approximately 40 epochs over the set of unique tokens.

We evaluated MutBERT and alternative models across 24 genomic datasets, with 23 assessed using the Matthews correlation coefficient (MCC) and one using the area under the receiver operating characteristic curve (AUROC). As shown in Fig. 2, MutBERT demonstrates strong performance across tasks assessed by the average MCC acorss 23 tasks, being only slightly outperformed by two models with significantly larger parameter counts. Additionally, we observed that MutBERT outperformed its two variant models (Fig. 2a). The two models that outperformed MutBERT were either substantially larger (NT-v1-2500M) or trained on significantly more diverse, multi-species datasets (NT-v2-500M). Models exclusively trained on human genome data, those trained on limited multi-species data, or those with moderate size (i.e., *≤*500M parameters) all underperformed compared to MutBERT. On the transcription factor binding site (TFBS) prediction task provided in GUE benchmarking, MutBERT series ranked as top performed models, with MuTBERT achieves the highest average MCC of 68.29, marginally surpassing DNABERT-2 (Fig. 2b). As shown in Fig. 2c, across the NT benchmark datasets, MutBERT achieves an average MCC of 65.82, outperforming DNABERT-2 and all Nucleotide Transformers trained exclusively on human data. It is only outperformed by Nucleotide Transformers with substantially more parameters and trained on more diverse datasets (e.g., NTv2- 500M-Multi and NT-2500M-Multi). However, we observed that MutBERT-Ref and MutBERT-Multi performed only comparably to other models. In Fig. 2d, for the eQTL variant effect prediction task, evaluated using average AUROC, MutBERT achieves a score of 0.615, slightly trailing MutBERT-Multi. Although models such as NT-v2-500M and NT-v1-2500M outperformed MutBERT, these models were trained either on extensive multi-species corpora or had substantially more parameters. As previously mentioned, MutBERT consistently ranked among the top-performing models trained exclusively on human genome data, limited multi-species datasets, or models with moderate parameter sizes. Overall, all three variants of MutBERT outperform many alternative models, particularly those with a comparable number of parameters. This highlights the effectiveness of our newly proposed model architecture, even when applied to models that take the reference genome sequence as input.

**Fig. 2:**
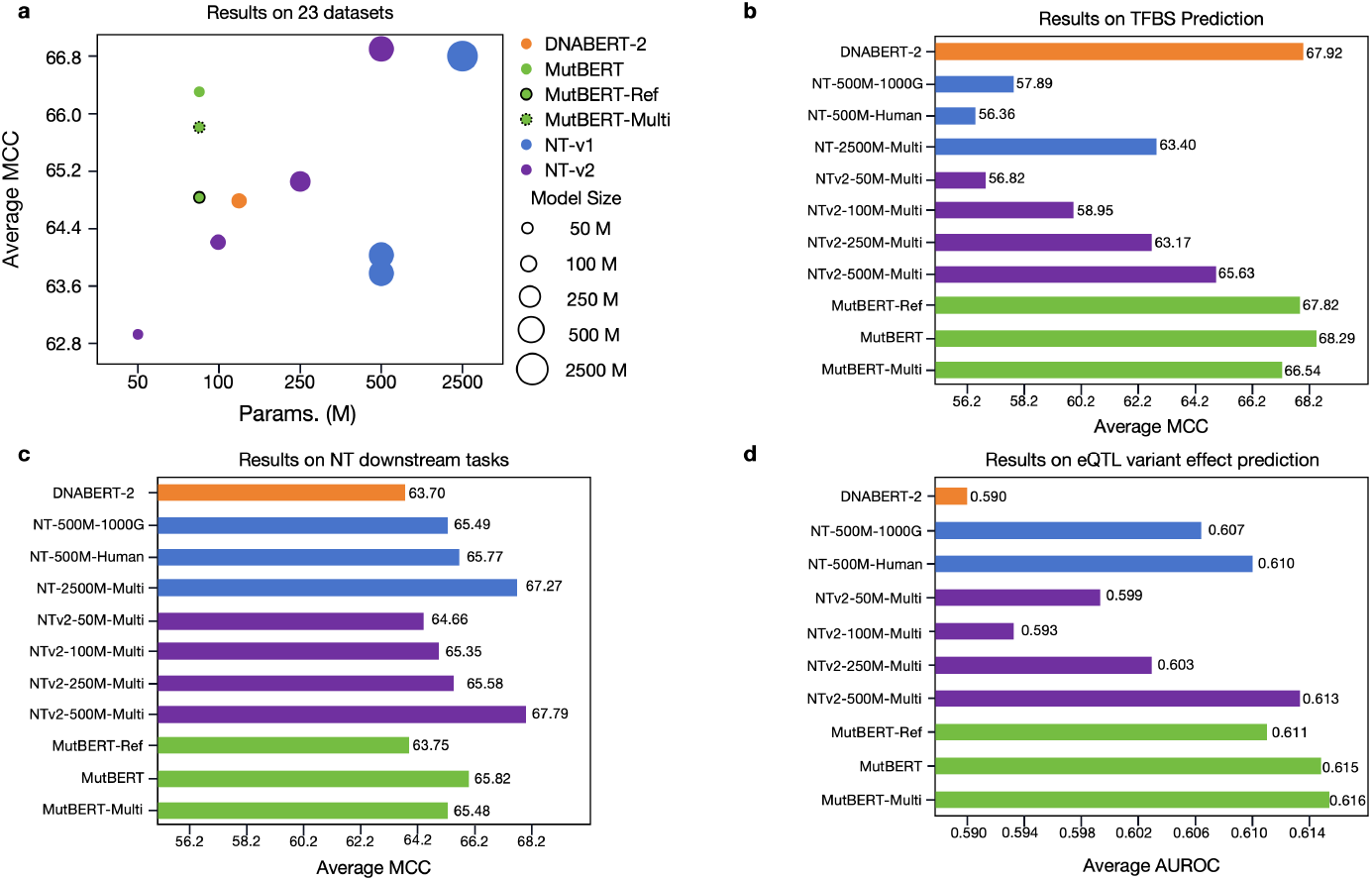
Results on multiple downstream tasks.

### TFBS Prediction and Nucleotide Transformer Tasks

Table 4 summarizes the performance of MutBERT and other models on the TFBS and Nucleotide Transformer downstream tasks, both evaluated using MCC. Larger models generally perform better, with NT-2500M-Multi consistently ranking among the top two models across most tasks. MutBERT ranked third in average MCC across 23 tasks, showcasing its strong generalization capacity despite its smaller size. Although MutBERT was the top-performing model in only 5 out of 23 individual tasks when considering all models, we observed that excluding models trained on substantially larger corpora (e.g., NT-v2) or models with significantly more parameters improved MutBERT’s relative performance. Under these conditions, MutBERT was the best-performing model in 13 out of 23 tasks.

**Table 4.** Comparison of MCC on TFBS and NT downstream task predictions. Tasks with ^*†*^ are from GUE benchmark, while the rest are from the NT benchmark. Models with * are fine-tuned with LoRA.

| Task | DNABERT-2 | MutBERT-Ref | MutBERT | MutBERT-Multi | NT-500M-1000G* | NT-500M-Human* | NTv2-50M-Multi | NTv2-100M-Multi | NTv2-250M-Multi | NTv2-500M-Multi* | NT-2500M-Multi* |
| --- | --- | --- | --- | --- | --- | --- | --- | --- | --- | --- | --- |
| TF-0<sup>†</sup> | 70.43 | **71.90** | 68.53 | <u>70.53</u> | 65.02 | 61.64 | 61.52 | 65.61 | 70.02 | 66.40 | 65.20 |
| TF-1<sup>†</sup> | 73.73 | **76.26** | <u>75.11</u> | 73.51 | 69.57 | 67.54 | 65.24 | 66.19 | 71.22 | 72.72 | 68.13 |
| TF-2<sup>†</sup> | **66.12** | 58.27 | 58.63 | 60.98 | 47.31 | 49.89 | 53.60 | 54.69 | 61.17 | <u>62.85</u> | 60.21 |
| TF-3<sup>†</sup> | 52.32 | **60.56** | <u>59.33</u> | 54.56 | 45.93 | 42.33 | 37.50 | 38.02 | 45.35 | 51.75 | 50.83 |
| TF-4<sup>†</sup> | <u>77.00</u> | 72.13 | **79.84** | 73.14 | 61.63 | 60.41 | 66.23 | 70.26 | 68.11 | 74.41 | 72.62 |
| H2AFZ | **52.79** | 50.65 | 49.61 | 50.00 | 49.86 | 48.78 | 49.15 | 50.50 | 48.91 | 51.10 | <u>52.75</u> |
| H3K27ac | 50.1 | <u>51.30</u> | 50.25 | 48.96 | 46.24 | 50.11 | 50.49 | 46.97 | 48.84 | 50.59 | **53.72** |
| H3K27me3 | 60.61 | 60.46 | **62.07** | 61.45 | 59.69 | 59.73 | 58.60 | 57.97 | 57.35 | <u>61.64</u> | 61.36 |
| H3K36me3 | 63.88 | 61.7 | 62.76 | 62.18 | 58.80 | 61.93 | 59.69 | 60.77 | 60.53 | <u>65.26</u> | **66.86** |
| H3K4me1 | 50.26 | 47.31 | 49.38 | 47.07 | 49.17 | <u>50.53</u> | 46.13 | 48.49 | 46.74 | **51.31** | 48.51 |
| H3K4me2 | <u>57.45</u> | 57.30 | 57.30 | 57.17 | 54.34 | 52.12 | 52.63 | 53.44 | 53.41 | **57.85** | 56.83 |
| H3K4me3 | 53.74 | <u>64.73</u> | 63.68 | 63.66 | 61.60 | 64.21 | 62.66 | 61.11 | 62.89 | **65.50** | 61.46 |
| H3K9ac | <u>55.18</u> | 51.9 | 52.06 | 52.97 | 50.94 | 52.22 | 50.42 | 45.55 | 53.64 | 54.96 | **57.89** |
| H3K9me3 | **52.24** | 48.78 | <u>50.60</u> | 48.94 | 44.95 | 46.06 | 41.04 | 44.76 | 38.24 | 48.02 | 47.14 |
| H4K20me1 | 64.33 | 65.42 | 65.78 | 66.7 | 62.01 | 65.62 | 64.82 | 62.65 | 65.28 | **67.74** | <u>67.46</u> |
| Enhancer | 54.31 | 52.47 | 51.26 | 51.9 | 57.18 | <u>59.02</u> | 53.01 | 56.57 | 55.13 | **60.19** | 57.85 |
| Enhancer types | 49.74 | 49.62 | 47.46 | 49.11 | <u>53.95</u> | 52.85 | 52.15 | 51.48 | 53.27 | **55.65** | 52.41 |
| Promoter all | 74.40 | 72.68 | 75.89 | 75.64 | 76.90 | 75.64 | 75.89 | 79.45 | 76.14 | **78.32** | <u>77.40</u> |
| Promoter non-TATA | 78.23 | 75.54 | 77.04 | 75.71 | 75.54 | 75.52 | 73.84 | 77.35 | 79.17 | <u>78.72</u> | 78.56 |
| Promoter TATA | 88.68 | 80.22 | 89.63 | 87.77 | **91.71** | 85.15 | 86.16 | <u>90.58</u> | <u>90.58</u> | 83.03 | 80.22 |
| Splice acceptor | 75.82 | 78.98 | 94.68 | 93.97 | 94.54 | 93.87 | 95.27 | 95.60 | **96.53** | **96.53** | <u>96.40</u> |
| Splice site all | 84.86 | 81.57 | 90.85 | 93.04 | 95.61 | 94.81 | 95.17 | 96.05 | <u>96.60</u> | 96.50 | **96.90** |
| Splice donor | 80.01 | 96.93 | 94.44 | 92.47 | 95.73 | 95.60 | 96.67 | 96.93 | <u>97.20</u> | **97.27** | 97.14 |
| Avg | 64.62 | 64.64 | <u>66.36</u> | 65.71 | 63.84 | 63.72 | 62.95 | 63.96 | 65.06 | **67.32** | <u>66.43</u> |

Further analysis revealed that MutBERT performs consistently well across all tasks, achieving results comparable to or slightly below the top-performing models. DNABERT-2 excels in TFBS and histone modification prediction tasks, while Nucleotide Transformers outperform in splice site prediction. Despite its compact size of 86 million parameters, MutBERT shows strong performance across both types of tasks, highlighting its versatility. Given that DNABERT-2 and Nucleotide Transformers primarily differ in tokenization methods, we hypothesize that BPE tokenization in DNABERT-2 is better suited for TFBS and histone modification predictions, whereas k-mer tokenization in NT is more effective for capturing individual base pair changes, as required for splice site prediction. Results from MutBERT-Ref suggest that single-nucleotide tokenization, as used in MutBERT, provides a balanced capability for both task types. Additionally, comparisons between MutBERT, MutBERT- Multi, and MutBERT-Ref highlight that the probabilistic genome representation used in MutBERT and MutBERT-Multi outperforms the deterministic reference genome sequence used in MutBERT-Ref in training genome language models. In addition, we observed a significant difference in performance on tasks related to splice-site prediction. NT models trained on extensive multi-species data handled these tasks much more effectively, significantly outperforming DNABERT-2 and slightly outperforming MutBERT and NT models trained primarily on human genome data. These results highlight the importance of further increasing the diversity of training data to enhance model performance.

### eQTL Variant Effect Prediction

We compared MutBERT with DNABERT-2 and Nucleotide Transformers on the eQTL variant effect prediction task, evaluated using AUROC. Due to GPU limitations, NT-2500M-Multi was excluded from this comparison. As shown in Table 5, MutBERT outperforms alternative models in most of the three distance categories, highlighting its strong mutation detection capabilities. Notably, for long-distance predictions, MutBERT benefits from the robust extrapolation ability of ROPE, which enables it to handle sequences significantly longer than those encountered during pretraining.

**Table 5.**
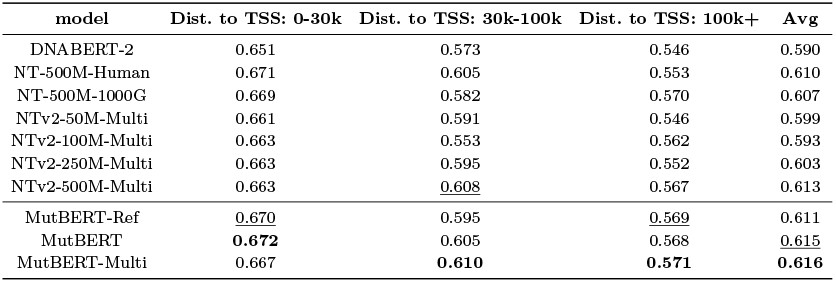
Comparison of AUROC on eQTL variant effect predicted by different models. The four columns represent distances to three different biological tissues and the average AUROC score.

| model | Dist. to TSS: 0-30k | Dist. to TSS: 30k-100k | Dist. to TSS: 100k+ | Avg |
| --- | --- | --- | --- | --- |
| DNABERT-2 | 0.651 | 0.573 | 0.546 | 0.590 |
| NT-500M-Human | 0.671 | 0.605 | 0.553 | 0.610 |
| NT-500M-1000G | 0.669 | 0.582 | 0.570 | 0.607 |
| NTv2-50M-Multi | 0.661 | 0.591 | 0.546 | 0.599 |
| NTv2-100M-Multi | 0.663 | 0.553 | 0.562 | 0.593 |
| NTv2-250M-Multi | 0.663 | 0.595 | 0.552 | 0.603 |
| NTv2-500M-Multi | 0.663 | <u>0.608</u> | 0.567 | 0.613 |
| MutBERT-Ref | <u>0.670</u> | 0.595 | <u>0.569</u> | 0.611 |
| MutBERT | **0.672** | 0.605 | 0.568 | <u>0.615</u> |
| MutBERT-Multi | 0.667 | **0.610** | **0.571** | **0.616** |

## Discussion and Conclusion

In this study, we introduced MutBERT, a probabilistic genome-based foundation model specifically designed to train genomic models on massive human genome data. Unlike previous approaches that often rely solely on individual genomes, MutBERT harnesses the diversity within population-scale genomic data by representing sequences as probabilistic distributions over allele frequencies. This approach allows MutBERT to directly learn variant information from a single probabilistic representation of the input sequence. In contrast, existing individual genome-based representations require models to learn from numerous separate genomes, repeatedly spanning the same SNV loci, which makes it challenging to effectively capture variant information due to their low occurrence.

Pre-trained exclusively on the human genome, MutBERT incorporates state-of-the-art techniques such as ROPE for efficient positional encoding, LoRA for parameter-efficient fine-tuning, and flash attention for computational optimization. We evaluated its performance across multiple downstream tasks using existing benchmarks, spanning 6 tasks and 24 datasets. Empirical results demonstrate that MutBERT consistently achieves competitive performance, securing top results in 5 of the 24 datasets and ranking among the leading models in the remaining benchmarks. When considering only models trained exclusively on human genomic data, this number increases to 13. These findings underscore MutBERT’s robustness and its ability to generalize across diverse genomic tasks, all while maintaining computational efficiency. MutBERT’s success stems from its ability to integrate population-level diversity into its training paradigm. For instance, consider a scenario where a model is trained on two human genomes. Traditional masked language modeling (MLM) methods would train the model to predict the most probable nucleotide at positions without single-nucleotide variants (SNVs) and assign high probabilities to observed alleles at positions with mutations. While effective, this approach often fails to fully utilize the richness of population-scale data. By contrast, MutBERT introduces a probabilistic framework that emphasizes allele frequencies, allowing the model to efficiently learn from population-level diversity without requiring explicit training on individual genomes. This representation strategy treats each genomic position independently, which works well in datasets like the 1000 Genomes Project, where SNVs are typically rare and spatially dispersed. In most cases, only one SNV appears within a 512- bp input window, making independent representation sufficient for capturing genomic variation. However, challenges arise when applying this framework to multi species datasets, where regions often contain high densities of mutations. In such cases, MutBERT’s independent representation may fail to capture co- variation across loci. Indeed, our results reveal that MutBERT’s performance on multi species datasets was lower than its performance on the 1000 Genomes Project data. Additionally, it remains unclear how MutBERT would perform if trained on probabilistic genomic sequences derived from larger and more diverse variant datasets, such as gnomAD (Karczewski et al., 2020), which exhibit denser SNV distributions. Although we followed common practice in population genetics by excluding rare variants from our analyses, this step is not strictly required in our framework. Thus, it will also be important for future studies to explore how including rare mutations affects MutBERT’s performance. All highlighting the need for future work to address these limitations.

Despite these limitations, MutBERT represents a significant advancement in genomic foundation modeling. Its probabilistic representation method enables efficient training and robust performance on population-scale genomic tasks. For example, we only pretrained our models on 4 L20 GPUs for 6 days. However, we recognize that further studies are needed to explore the potential of scaling up MutBERT. We hypothesize that increasing its model size could further enhance its performance, potentially surpassing larger models while retaining its computational efficiency.

In conclusion, MutBERT sets a new standard for genomic representation learning by bridging the gap between computational efficiency and biological insight. Its innovations provide a strong foundation for leveraging biobank-scale genomic data.

## Author contributions

W.L. and Y.Z. conceived the study. W.L., H.S., and J.X., performed analysis. Y.Z. supervised the project. W.L. and Y.Z. wrote the article. All authors read and approved the final article.

## Acknowledgements

We thanks DNABERT-2’s authors for their helps.

